# Multidomain Analysis of Clinical Cognitive Assessments and Imaging Data in Alzheimer’s Disease Accurately Predicts Disease Stage and Grade Independent of Amyloid and Tau

**DOI:** 10.64898/2026.04.12.717232

**Authors:** Juan Antonio K. Chong Chie, Scott A. Persohn, Olivia R. Simcox, Paul Salama, Paul R. Territo, the Alzheimer’s Disease Neuroimaging Initiative

**Author notes:** Data used in preparation of this article were obtained from the Alzheimer’s Disease Neuroimaging Initiative (ADNI) database (adni.loni.usc.edu). As such, the investigators within the ADNI contributed to the design and implementation of ADNI and/or provided data but did not participate in analysis or writing of this report. A complete listing of ADNI investigators can be found at: http://adni.loni.usc.edu/wp-content/uploads/how_to_apply/ADNI_Acknowledgement_List.pdf.

## Abstract

**Background:** Individual clinical cognitive assessments (CCA) for Alzheimer’s disease (AD) provide broad disease stratification but are limited in sensitivity and specificity, requiring integration of multiple CCA for optimal disease staging. Recent work from our lab suggests that neuro-metabolic and vascular dysregulation (MVD) occurs early in AD, prior to clinical symptoms, and may provide higher sensitivity and specificity than CCA alone. In this study, we combined three widely accepted CCA with MVD readouts and developed a multimodal ensemble machine learning approach across the AD spectrum to predict disease stage and grade.

**Methods:** AD subjects (N=372) across the disease spectrum with imaging (PET:18F-FDG, MRI:T1w, T2 FLAIR, ASL) and CCAs (ADAS-Cog, CDR, MoCA) data were analyzed from ADNI. Imaging data were registered to MNI152+, z-scored relative to cognitively normal controls, and processed for MVD. A clinical-set-enrichment analysis (CSEA) was developed to link regional brain changes with CCA scores, map changes to functional categories, project them into a 3D Cartesian space, and model trajectories, thus revealing at-risk and resilient regions. In addition, an ensemble machine-learning approach was utilized for disease stage classification, and a disease grading scheme across the AD spectrum was developed to further stratify within disease stages.

**Findings:** Regional data followed an MVD pattern across AD stages stratified by CSEA scores. Females showed greater stage separation along the CCA axis within each region, indicating faster disease progression. Moreover, progression in at-risk brain regions (e.g., mid- and inf-temporal gyri, amygdala) was associated with longer disease path lengths, whereas progression in resilient brain regions (supramarginal gyrus) was not. Moreover, our classification and grading approach can predict AD stage and grade independent of amyloid-beta and tau with high precision and accuracy.

**Interpretation:** A framework was developed to evaluate MVD and CCA variations across the AD spectrum, thereby distinguishing at-risk and resilient brain regions. Distinct disease trajectories were identified, and a new data-driven grading scheme was proposed to highlight the potential for precision medicine and therapeutic evaluation.

**Funding:** NIH T32AG071444

## 1. Introduction

Alzheimer’s Disease (AD) is a neurodegenerative disorder characterized by cognitive decline, language dysfunction, motor impairment, and visuospatial deficits.^1^ Clinically, physicians evaluate these changes in patients using a variety of clinical cognitive assessments (CCAs). However, these assessments often have broad stratifications, reduced sensitivity due to the overlap between cognitive domains, and may not accurately reflect real-world function.^2^ In addition, these assessments may not capture the early stages of AD, when clinical manifestations are not yet evident, thereby affecting their accuracy.^1^ Among CCAs, common screening tests include the Alzheimer’s Disease Assessment Scale-Cognitive Subscale (ADAS-Cog),^3^ Clinical Dementia Rating (CDR),^4^ and Montreal Cognitive Assessment (MoCA).^5^ While these assessments offer the advantage of being performed quickly and in primary care, they also have significant limitations, including cultural and language biases, cut-off scores that can lead to false positives or negatives, and the potential to miss early, subtle cognitive decline.^6-9^ Currently, no single method provides a comprehensive assessment; thus, multiple assessments must be integrated to achieve optimal assessment.^9-11^

Recent studies conducted in preclinical and clinical settings to evaluate neuro-metabolic and vascular dysregulation (MVD) suggest that this dysregulation occurs early in the disease spectrum, even before the manifestation of clinical symptoms, and prior to the accumulation of amyloid-β (Aβ) and tau tangles.^12-15^ These studies have significantly contributed to our understanding of disease etiology by proposing several theories of the changes the brain undergoes across the disease spectrum.^16-21^ For instance, several lines of evidence suggest that gliosis is a response mechanism to oxidative stress and to alterations in cerebral metabolism and perfusion.^22,23^ This response triggers a series of events that result in transcriptomic alterations (e.g., mitochondrial and lipid metabolism changes, oxidative stress, vascular alterations),^24^ microglial activation,^25^ and astrocytosis,^26,27^ which occurs prior to cognitive decline.^26,28^ Conversely, the metabolic reprogramming theory^17,18^ proposes that metabolic dysregulation in neurons and astrocytes leads to bioenergetic deficits, contributing to neuronal degeneration and ultimately causing AD onset.^17,18^ A recent study conducted by our lab suggests that these changes can be quantified, and used to identify inter-regional functional alterations via Region-Set Enrichment Analysis (RSEA).^29^

In addition to MVD and bioenergetic changes, several studies revealed that brain regions progress at different rates.^13,30,31^ Regions such as the hippocampus, mid-temporal gyrus, amygdala, and para-hippocampal gyrus are more susceptible,^13,30^ and show substantial atrophy in prodromal AD stages.^31^ Moreover, Aβ and tau deposition follow a regionally specific and sequential pattern, further supporting this theory.^32,33^ Consequently, early MVD alterations alone may not be accurately measured using CCA evaluations, since patients do not manifest clinical symptoms at this stage.^13,34^ Moreover, since these tests lack the ability to evaluate brain-region-level changes, unlike imaging biomarkers, this suggests that a combination of biomarkers is needed to understand disease progression, especially at early stages.^35^

Recent advances in machine learning and artificial intelligence approaches (ML/AI) have been leveraged for tasks such as disease classification,^35^ prognosis prediction,^35^ and neuroimaging segmentation.^36^ However, the quality and type of input data directly impact the performance and reliability of ML/AI approaches.^37^ Despite the growing emphasis on explainable and interpretable ML/AI, the typical “black-box” nature of many of these methods hinders interpretability, limiting their utility for explaining underlying disease mechanisms.^38^ Moreover, the limited generalizability across populations and the systematic biases introduced by imbalanced training datasets pose challenges to the deployment of ML/AI in clinical practice.^39^

In this study, expanding on our previous work,^13,29^ we developed a new method to combine three widely accepted CCA evaluations (ADAS-Cog, CDR, MoCA) into a composite measure aligned with brain regions. Provided this, we explored a multidomain approach to evaluate the relationship between MVD imaging biomarkers and CCA measures, resulting in a new cognitive degradation surrogate index. These combined measures aim to leverage the strengths of each domain into a single readout, thus improving the sensitivity and specificity of the independent readouts across the disease spectrum. Therefore, we hypothesize that combining these biomarkers with vitals (e.g., age, BMI, sex) and genetics (e.g., *APOE*) can further facilitate disease stratification, yielding a finer gradation across the AD spectrum.

## 2. Methods

### 2.1 Dataset

Data used in the preparation of this article were obtained from the Alzheimer’s Disease Neuroimaging Initiative (ADNI) database (adni.loni.usc.edu). ADNI was launched in 2003 as a public-private partnership, led by Principal Investigator Michael W. Weiner, MD. The primary goal of ADNI has been to test whether serial magnetic resonance imaging (MRI), positron emission tomography (PET), other biological markers, and clinical and neuropsychological assessment can be combined to measure the progression of mild cognitive impairment (MCI) and early AD.^40^

The dataset consists of 372 clinical subjects from ADNI phases 2 and 3 that have all the following data: vitals measurements (age, sex, height, weight), *Apolipoprotein E* (*APOE)* genotype, ^18^F-FDG PET, T1-weighted (T1w), T2 Fluid-Attenuated Inversion Recovery (T2-FLAIR), Arterial Spin Labeling (ASL) MRI, ADAS-Cog, CDR, and MoCA within 180 days from each other. Images were obtained according to standardized protocols developed by ADNI,^40^ and were reconstructed using standard parameters (Table S1-S2). From this population, subjects were aged 55-95 years, and 52.2% were male. The split between disease stages was: cognitive normal (CN)=84, early-MCI (EMCI)=70, MCI=103, late-MCI (LMCI)=40, and AD=75. To ensure uniformity across CCA scales, MoCA scores were rescaled to match the ascending score pattern followed by ADAS-Cog and CDR. For a subpopulation, cerebrospinal fluid (CSF) biomarkers (Aβ42, p-Tau-181, t-Tau) were mined, with a split of: CN=64, EMCI=56, MCI=71, LMCI=39, and AD=66.

For the ML/AI sections, the dataset was divided into two sets: the first set consists of 80% of the data used for training and validation of the approach, while the second set consists of the remaining 20% used for testing to evaluate the performance of the ML/AI approach for unseen data.

### 2.2 Imaging Biomarkers

^18^F-FDG PET was used as a surrogate marker of glucose uptake to evaluate regional cerebral glycolysis.^41^ T1w, T2-FLAIR, and ASL MRI were used to generate cerebral blood flow (CBF) maps via ExploreASL.^42^ To allow a direct comparison at the population level, images were co-registered to a common reference space (Montreal Neurological Institute + Harvard-Oxford Cortical + subcortical (RRID: SCR_001476) + FSL Probabilistic Cerebellar Atlases; MNI152+). Mean intensities for 59 brain regions were extracted via the MNI152+ for ^18^F-FDG and CBF maps. Per our previous studies,^13,29^ regional mean intensity values for each imaging modality were ratioed relative to the whole-brain value to obtain standardized uptake value ratios (SUVR), which were then converted to z-scores relative to the CN population.

### 2.3 Clinical-Set Enrichment and Functional-Set Enrichment Analyses

Following a similar process to a previous study in our lab,^29^ clinical-set enrichment analysis (CSEA) was developed to quantify brain functional changes based on CCA evaluations. Although this method was developed using three different CCAs (ADAS-Cog, CDR, MoCA), it can be expanded to accommodate different combinations and types of CCAs. This approach maps CCA evaluations into 10 functional categories (FC): auditory, autonomics, cognition, emotions, language, memory, motor, sensorial, speech, and visual. Therefore, an *m* × 10 matrix is generated for each CCA evaluation, where the number of rows (*m*) is the number of tasks or elements in the CCA evaluation (13 for ADAS-Cog, 6 for CDR, 8 for MoCA), and the number of columns represents the different FCs. Due to the lack of a surrogate measure for autonomics functions, this category was excluded from the analysis, and the column was set to 0 in subsequent steps. To populate these matrices, a column is assigned a value of 1 for each CCA evaluation if the CCA task aims to measure the corresponding FC; otherwise, it is assigned a value of 0. Then, each value is normalized using the following formula:

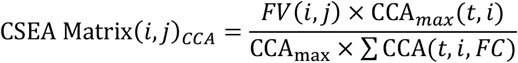

where *i* is the CCA task index, *j* is the FC index, *t* is the task, and *FV*(*i, j*) is the entry (*i, j*) of the *m* × 10 CSEA_CCA_ matrix. Then, the CSEA_CCA_ matrices are normalized by dividing each CSEA_CCA_ matrix by the number of assessments (*n*_*a*_) used (in this study *n*_*a*_ = 3) and subsequently concatenated to create a single CSEA matrix. In this study, the final CSEA matrix is of size 27 × 10.

To perform CSEA analysis, CCA scores for each task are concatenated and multiplied element-wise across the corresponding CSEA matrix row; this process generates a CSEA Score Matrix (CSM), where each row represents the changes measured for each task across the 10 different FCs. Furthermore, these changes can be mapped to brain regional changes using RSEA, as developed in a previous work from our lab,^29^ by multiplying the CSEA scores by the transpose of the RSEA matrix, a process called functional-set enrichment analysis (FSEA):

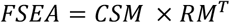

where FSEA is a matrix of size 27 × 59 for the 27 different CCA tasks across the 59 brain regions in the MNI152+, and RM is the RSEA matrix of size 59 × 10 for 59 brain regions in the MNI152+ atlas and the 10 FCs. This process allows mapping of changes in the CCA score by brain region. However, as mentioned before, due to a lack of a precise surrogate measure for autonomics functions, the four brain regions (Central Opercular Cortex, Cerebral White Matter, Lateral Ventricle, Brain Stem) related to this category were excluded from the analysis and were set to 0, reducing the number of regions to 55. Lastly, the FC score for each brain region was generated by summing the columns of the FSEA matrix, yielding a 1 × 59 regional cognition index (rCI) vector.

### 2.4 Multidimensional Analysis

To permit tracking of the trajectories for individual brain regions across the disease spectrum, z-scored regional SURV values for glycolytic metabolism (rCGM), z-scored regional SURV values for cerebral perfusion (rCBF), and rCI for each brain region were plotted into the *x, y*, and *z* axes of a 3D Cartesian space, respectively (Fig.1). In this 3D space, metabolic and perfusion measures span the interval (−∞, ∞), cognition indices span the interval [0, ∞), and the origin of this 3D space (location (0,0,0)) represents the CN population. As described in our previous work,^13^ z-scored metabolic and perfusion measurements oscillate between downregulation and upregulation across the AD spectrum; thus, it is a necessary to allow these values to span this interval. On the other hand, CCA assessment usually assigns a fixed score to CN subjects (0 for ADAS-Cog and CDR, 30 for MoCA), and then this score is mapped onto a scale to indicate cognitive decline; thus, negative values for cognition indices lack “real-world” meaning.

**Figure 1.**
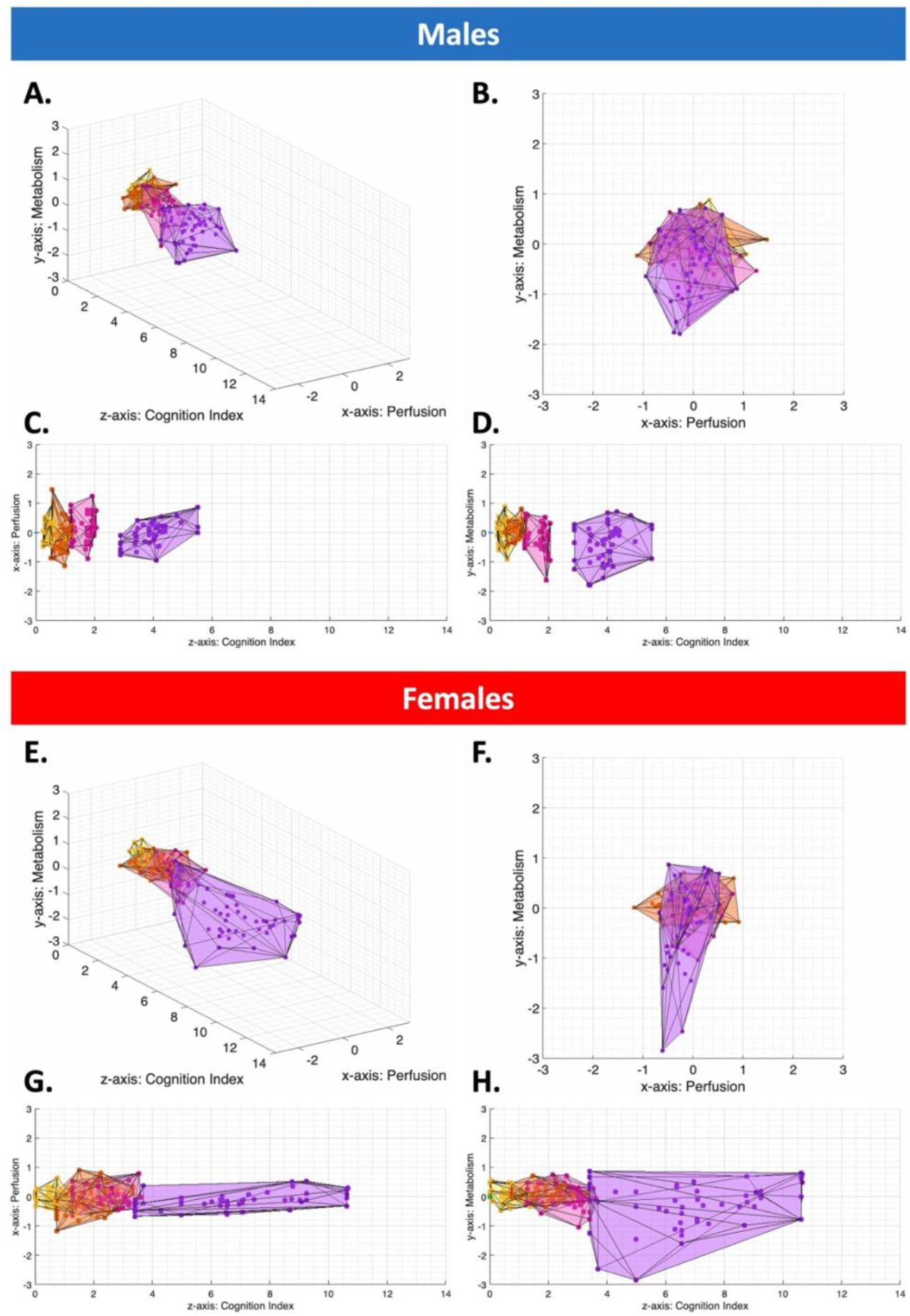
3D Multi-Domain Space Plots. Plots for the 3D projections of z-scored cerebral perfusion (x-axis), z-score cerebral metabolism (y-axis), and regional cognitive index (z-axis), where each dot represents the cross-population average for the respective disease stage. (A-D) (E-H) are the 3D multi-domain results for the male subjects, and (A-D) are the 3D multi-domain results for the female subjects. (A,E) represent the 3D view, (B,F) represent the 2D view for the x-y plane, (C,G) represent the 2D view for the x-z plane, and (D,H) represent the 2D view for the y-z plane. Males show more compact convex hulls; in contrast, females are more widely distributed across the cognition index, highlighting sex-dependent differences.

Regional brain analysis used rCGM, rCBF, and rCI values across disease stages as spatial 3D coordinates. Typically, neurodegeneration develops gradually over extended periods, reflecting progressive neuronal dysfunction and loss driven by the accumulation of pathological changes.^43,44^ In addition, the MVD pattern does not follow a linear progression across the disease spectrum.^13,14^ Hence, to characterize the individual regional trajectories across the 3D multi-domain disease spectrum, 3D cubic-spline fitting was selected because it is an interpolation technique designed to fit continuous curves while minimizing abrupt changes by enforcing smoothness constraints, without sacrificing flexibility.^45,46^ Cubic-spline interpolation was performed as described by de Boor,^47^ using cross-population disease-stage coordinates to evaluate the trajectories followed by each brain region.

### 2.5 Machine Learning Disease Classification

In ML/AI, ensemble methods combine multiple models (or learners) to achieve better predictive performance than a single model.^48,49^ Boosting is a type of ensemble approach in which learners are trained sequentially, with each new learner focusing more on training instances that previous learners misclassified by increasing their weights, thereby reducing bias and improving predictive accuracy and generalization.^48^

In this study, to evaluate the potential and accuracy of ML/AI methods to accurately classify and stage subjects based on multi-domain readouts, we used Adaptive Boosting (AdaBoost),^50,51^ a boosting-based ensemble algorithm that creates a strong classifier by combining multiple ‘weak learners’ (WL) whose individual performance is typically only slightly better than random guessing. At each iteration, AdaBoost updates the training data weights by assigning higher weights to misclassified instances, then estimates a coefficient for the new WL based on its accuracy. The final prediction is obtained by a weighted sum over all WLs, with more accurate learners having greater influence. Through this adaptive reweighting of both the training data and WL, AdaBoost typically converts a collection of WLs into a strong model that generalizes better to unseen data. Fig.2.A shows the descriptive diagram of this model.

**Figure 2.**
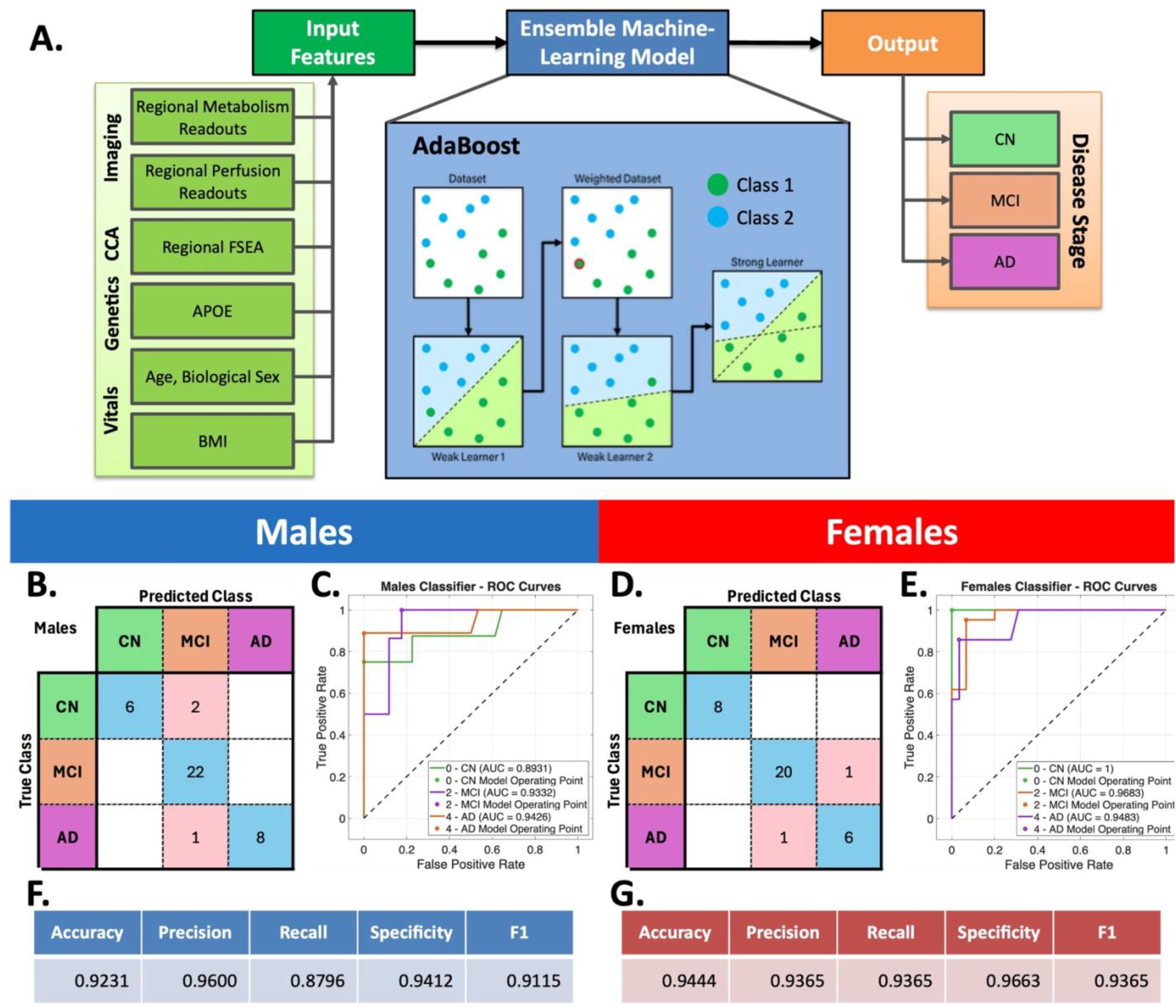
Machine Learning Framework and Results. (A) Schematic representation of the implemented machine learning framework for AD stage classification. (B) Confusion matrix for the male subjects in the testing set, with the main diagonal representing correctly classified cases and off-diagonal elements representing misclassifications. (C) ROC curves for the male subjects. (D) Confusion matrix for the female subjects in the testing set. (E) ROC curves for the female subjects. (F,G) Standard classification performance matrices for males and females, respectively. Overall, the framework misclassified only 5 out of 75 subjects in the testing set. These findings demonstrate that the disease classification framework achieves high performance across multiple evaluation metrics.

Given the sex-dimorphism effects observed in our previous works,^13,29^ two models were stratified by sex and trained using a 5-fold cross-validation scheme. The input features for each model, per subject, were: age, body-mass index (BMI), *APOE* genotype, and 3D multi-domain values (rCGM, rCBF, rCI). AdaBoost hyperparameters (e.g., learning rate, number of learning cycles, number of WL) were optimized using Bayesian hyperparameter optimization, a strategy that uses a probabilistic surrogate model and Bayes’ theorem to efficiently explore the hyperparameter space.^52-54^ Model performance was measured using receiver operating characteristic (ROC) curves, the area under the ROC curve (AUC), and classification performance metrics (accuracy, precision, recall, specificity, F1-score).

### 2.6 Disease Grading Scheme

In this study, we developed a disease grading scheme (Fig.3.A) to further stratify subjects within a disease stage using the regional trajectories observed in the 3D multidomain space (Fig.3.B). For each regional cubic spline (Fig.3.C), we developed a method called <u>U</u>nwrapped <u>N</u>ode <u>F</u>unction for <u>O</u>ptimized <u>L</u>ength <u>D</u>etermination (UNFOLD), which linearizes the spline and permits quantization of the changes on a linear scale (Fig.3.D). This method works by calculating the spline length between each disease stage to determine the location of the cross-population averages over the interval [0, ∞). For details, see Supplemental Methods A.1.

**Figure 3.**
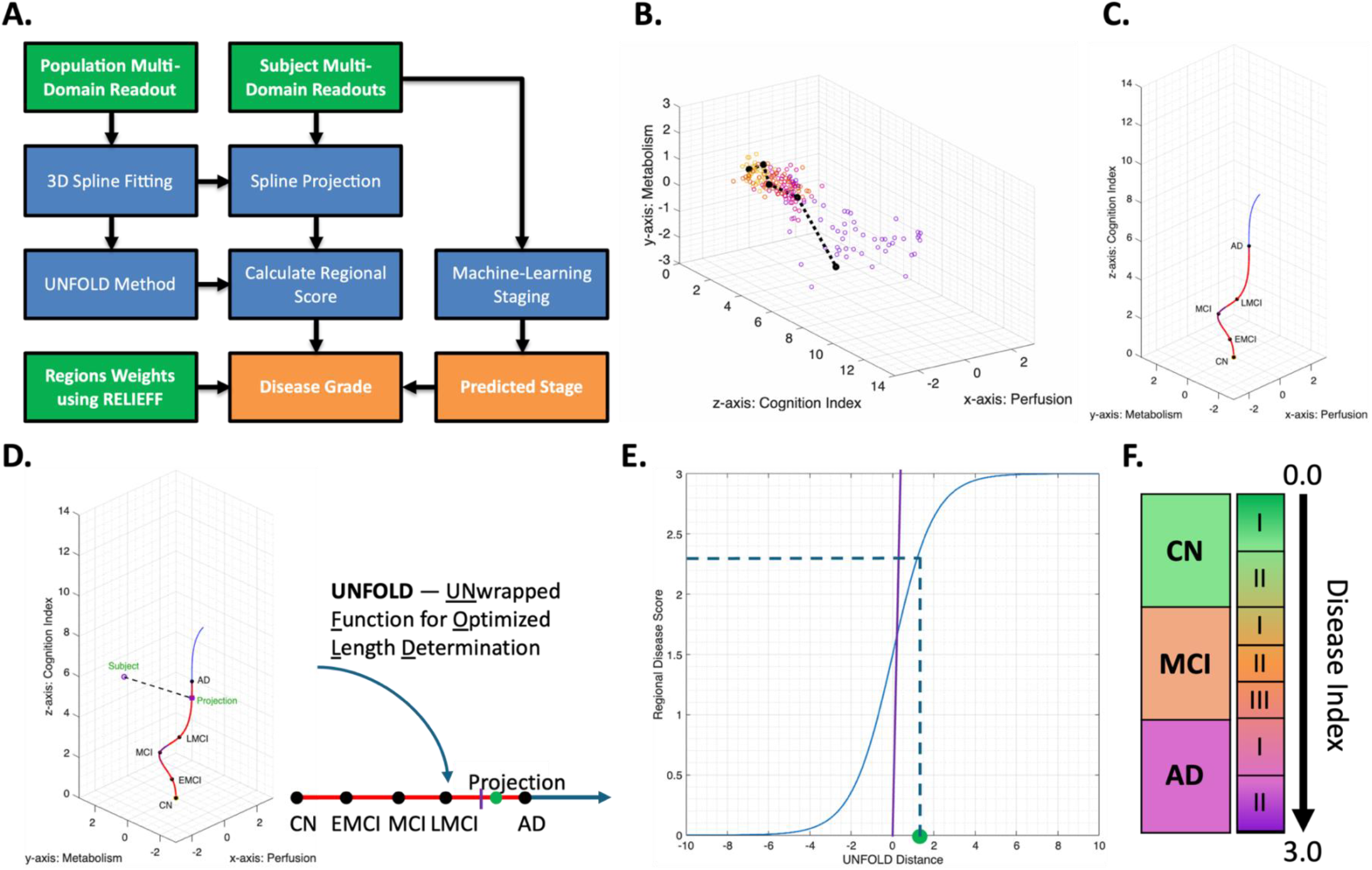
Stage Grading Framework. (A) Schematic representation of the AD stage grading scheme using 3D multi-domain readouts. (B) Trajectory of region #13, the mid-temporal gyrus temporooccipital, within the 3D multi-domain space. (C) 3D cubic spline fitted to the trajectory of region #13. (D) Illustration of the Unwrapped Function for Optimized Length Determination (UNFOLD) process and projection of an individual subject onto the 3D cubic spline. (E) Regional disease score calculated based on the UNFOLD-projected distance. (F) Proposed grading scale for further stratification within the AD disease stage. Steps (B-E) are performed for each region, and the final disease index is computed as the weighted sum of the regional disease scores.

UNFOLD distance to the disease index score, were fitted with a sigmoid function, which is consistent with widely used dynamic models of AD biomarker trajectories, and has been shown in several studies to follow non-linear sigmoidal distributions across the disease continuum,^55-57^ where the data range for the disease index were bound over the interval [0,1]. Then, for the disease stages CN and AD, a sigmoid distribution is fit under the condition that the UNFOLD midpoint distance between the two consecutive stages (CN with EMCI and LMCI with AD) is 0.5. For the set of disease stages EMCI, MCI, and LMCI, the midpoint between EMCI-MCI and MCI-LMCI are set to a quarter log-steps below and above from the midpoint of the sigmoid, respectively. In the case of CN and AD, by constraining the midpoint of the sigmoid at 0.5, we enforce that the UNFOLD distance between the two stages is mapped to the center of the grading scale, which makes the grading symmetric with respect to both stages. Similarly, for the EMCI-MCI-LMCI block, setting the cuts at 1/3 and 2/3 partitions the [0,1] interval into three uniform, ordinally spaced subintervals.

To generate the grade for a subject, each region is projected into the spline to find its 3D location, which is subsequently projected to the UNFOLD linearized scale (Fig.3.D). The regional disease score is computed by finding its value in the sigmoid distribution corresponding to the UNFOLD location (Fig.3.E). To combine regional disease scores into a single subject score, the regional scores are multiplied by normalized regional weights computed using ReliefF,^58^ a feature selection algorithm that assigns an importance ranking and weight to each feature. Lastly, the weighted regional values are summed to generate a single value. This value spans over the interval [0,3], where the intervals [0,0.5), [0.5, 1), [1, 1.33), [1.33, 1.67), [1.67, 2), [2,2.5), and[2.5,3] correspond to the grades CN I, CN II, MCI I, MCI II, MCI III, AD I, and AD II, which is illustrated in Fig.3.F.

### 2.7 Statistical Significance and Performance Evaluation

The goodness of fit between the regional 3D Cartesian coordinates and the regional 3D cubic splines was evaluated using R-squared. Classification model performance was measured using AUC for ROC curves and standard classification performance metrics (e.g., accuracy, specificity). Feature ranking and feature weights were measured using ReliefF.^58^ Correlation between disease index score and CCA was quantified using Pearson’s correlation coefficient, and its significance was evaluated with a t-statistic under the null hypothesis of zero correlation.

## 3. Results

### 3.1 Multidimensional Analysis of Cerebral Metabolism, Cerebral Perfusion, and Clinical Assessments

Our lab has developed a framework to evaluate MVD and CCA variations by projecting surrogate readouts of cerebral metabolism and perfusion, combined with CCA evaluations at a brain regional level, onto a 3D Cartesian space (Fig.1). The *x, y*, and *z* axes represent changes in glycolytic metabolism, cerebral blood flow, and CCA relative to the CN population, respectively. Using this framework, we observed that regional metabolism and perfusion changes follow an MVD pattern, consistent with our previous research. ^13^ Importantly, the addition of the rCI as a third-axis representing cognitive decline based on the CCA evaluations further stratifies these changes. Notably, the four disease stages (EMCI, MCI, LMCI, AD) relative to CN can be clustered and follow a distinct disease trajectory (Fig.1). In addition, males (Fig.1.A-D) and males (Fig.1.E-H) follow similar trends, with females exhibiting a broader spread across the cognitive index axis and males’ clusters being more compact, as measured by the isoperimetric quotient^59^ for males/females: EMCI=0.5115/0.6104, MCI=0.5092/0.4114, LMCI=0.5007/0.4902, AD=0.5459/0.15260, where larger values imply more compact figures.

The regional trajectories showed that each region progressed at a different pace through the disease stages, as assessed by the total length of the vectors connecting two consecutive stages (Fig.3.B-C). For example, regions such as the mid-temporal gyrus, amygdala, and subcallosal cortex have longer disease path length than regions such as the supramarginal gyrus, consistent with the hypothesis that path lengths indicate at-risk or resilient status of each brain region across the AD continuum (Fig.S1). Since the length of the path is related to the stage and the MVD trajectory, long path lengths are observed in regions known to be at risk, while short path lengths are observed in brain regions known to progress more slowly in AD and represent a resilience phenotype. In addition, since these brain regions’ coordinates indicate 3D spatial localization at the population level, connecting them with a straight line between segments may underestimate the actual trajectory length. Notably, sex differences are observed, with most brain regions in AD showing higher cognitive index values in females. This suggests faster progression and/or more pronounced clinical manifestations.

To approximate these trajectories, cubic splines were fitted to the 3D coordinates of each disease stage, and all splines started at the origin. For the mid temporal gyrus, the R-squared between the fitted cubic splines and the linear approximation is 0.9930/0.9946 for males/females (Fig.3.C), and across brain regions the average R-squared is 0.9926±0.0096/0.9876±0.0095 for males/females. For individual regional R-squared, see Table S3. Modeling the multi-domain splines reveals that regions follow a spirally curved trajectory, with changes in curvature dependent on subject sex, brain region resilience, and disease stage. Although many brain regions follow the spiral curve (Fig.4.A), some regions do not strictly follow this pattern, such as the para-hippocampal gyrus (Fig.4.B-C), suggesting distinct trends between regions which could be due to resilience status, MVD dysregulation limits, region-specific susceptibility to AD pathology, or involvement of co-pathologies.

**Figure 4.**
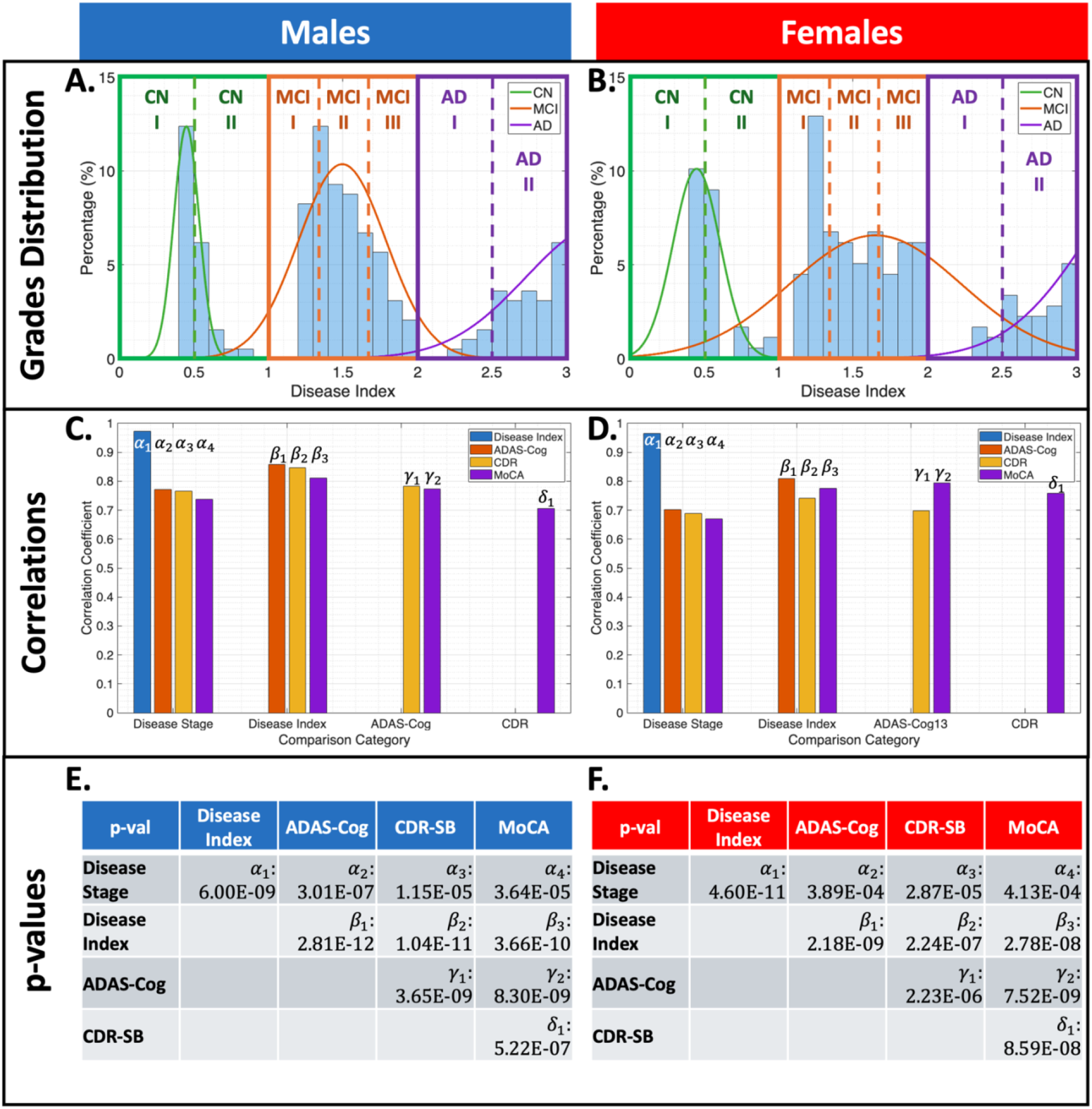
Results of the Stage Grading Scheme. ((A,B) Histograms of grades and disease index for (A) males and (B) females, overlaid with fitted Gaussian distributions for CN (green), MCI (orange), and AD (purple). Dotted lines represent grade boundaries. (C,D) Correlation coefficients between disease stage, disease index, and cognitive clinical assessments (ADAS-Cog, CDR-SB, MoCA score) for (C) males and (D) females. (E,F) Pairwise p-values for (E) males and (F) females indicate that all results were statistically significant, supporting disease index as a robust marker of disease severity.

### 3.2 Machine Learning Disease Classification Results

By analyzing cross-sectional population averages in the 3D multi-domain space (Fig.1), it can be observed that each disease-stage region cloud is roughly separable, suggesting that a machine learning model can classify and stage subjects from these readouts with high accuracy. To test and evaluate the feasibility of this hypothesis, an AdaBoost model was trained on 80% (N=297) of the data and subsequently tested on the remaining 20% (N=75) as described in Fig.2.A. This model achieved an overall AUC of 0.9476 and a classification accuracy of 93.33% (males=92.31%, females=94.44%) with only 5 misclassified cases out of 75 (3/39 for males and 2/36 for females). Performance of the model is shown in Fig.2.B-G

Consistent with the confusion matrices, the model achieved high precision (males=0.9600, females=0.9365, overall=0.9488), indicating that when the classifier assigns a subject to a given clinical stage, it is correct most of the time with few false positives. The specificity was also high (males=0.9412, females=0.9663, overall=0.9527), indicating that the classifier reliably identifies non-cases for each class and rarely mislabels CN subjects in females or MCI subjects in males. Misclassified cases occurred mainly at the boundaries between adjacent stages (e.g., CN-vs-MCI, MCI-vs-AD), consistent with the biological and clinical overlap between these groups and suggesting that residual errors largely reflect transitional phenotypes rather than gross mislabeling.

### 3.3 Disease Grading Results

To evaluate the distribution of the disease index across disease stages, the disease index values were computed for the complete dataset (N=372), using the pipeline shown in Fig.3.A. Each stage exhibits a pattern that closely resembles a Gaussian distribution (Fig.4.A-B), indicating a consistent grading profile within groups. By overlaying the different disease stage grades, the distribution shows an evident separation and a progressive shift across stage grades, illustrating how the grades from successive disease stages are distributed along the overall AD continuum. This visualization highlights both within-stage variability and the gradual shift in the grade distribution across disease progression.

Next, we assessed the relationship between the disease index and each CCA using Pearson’s correlation coefficient. The disease index showed a significantly stronger correlation with disease stage than any individual CCA, suggesting that it better reflects overall disease severity (Fig.4.C-F). In addition, the disease index was more significantly correlated with the CCAs than the CCAs were with one another, highlighting its ability to integrate information from multiple data domains (e.g., imaging, clinical assessments), which better represents the underlying disease continuum. These results are shown in Fig.4.C-D, and the p-values of the correlations are shown in Fig.4.E-F.

Last, we examined the relationship between our disease index values and CSF biomarker levels for Aβ and tau accumulation (Fig.5). Notably, the disease staging and grading framework assigns stage and grade independent of Aβ and tau readouts. The available data from the ADNI subjects were CSF values for Aβ42 (pg/ml), p-Tau-181 (pg/ml), and t-Tau-181 (pg/ml). Using this data, we derived Amyloid±/Tau± profiles using the p-Tau-181 ratio with AB42 and the p-Tau-181 value, with cutoff values previously reported by van Harten.^60^ Across disease index grades, CSF amyloid and tau biomarkers showed a clear shift from predominantly negative to predominantly positive profiles in both males and females. Subjects in the early grades (CN I, CN II, and MCI I) were predominantly amyloid-negative and had low p-tau-181 values, suggesting limited biomarker evidence of AD despite varying disease indices. At MCI II and MCI III, a subset of subjects became amyloid- and tau-positive, suggesting that the disease index captures emerging pathological burden before it becomes universal. In the highest grades classified as AD I and AD II, nearly all individuals were amyloid-positive and showed elevated p-tau concentrations, consistent with an advanced biomarker-defined Alzheimer’s pathophysiologic continuum.

**Figure 5.**
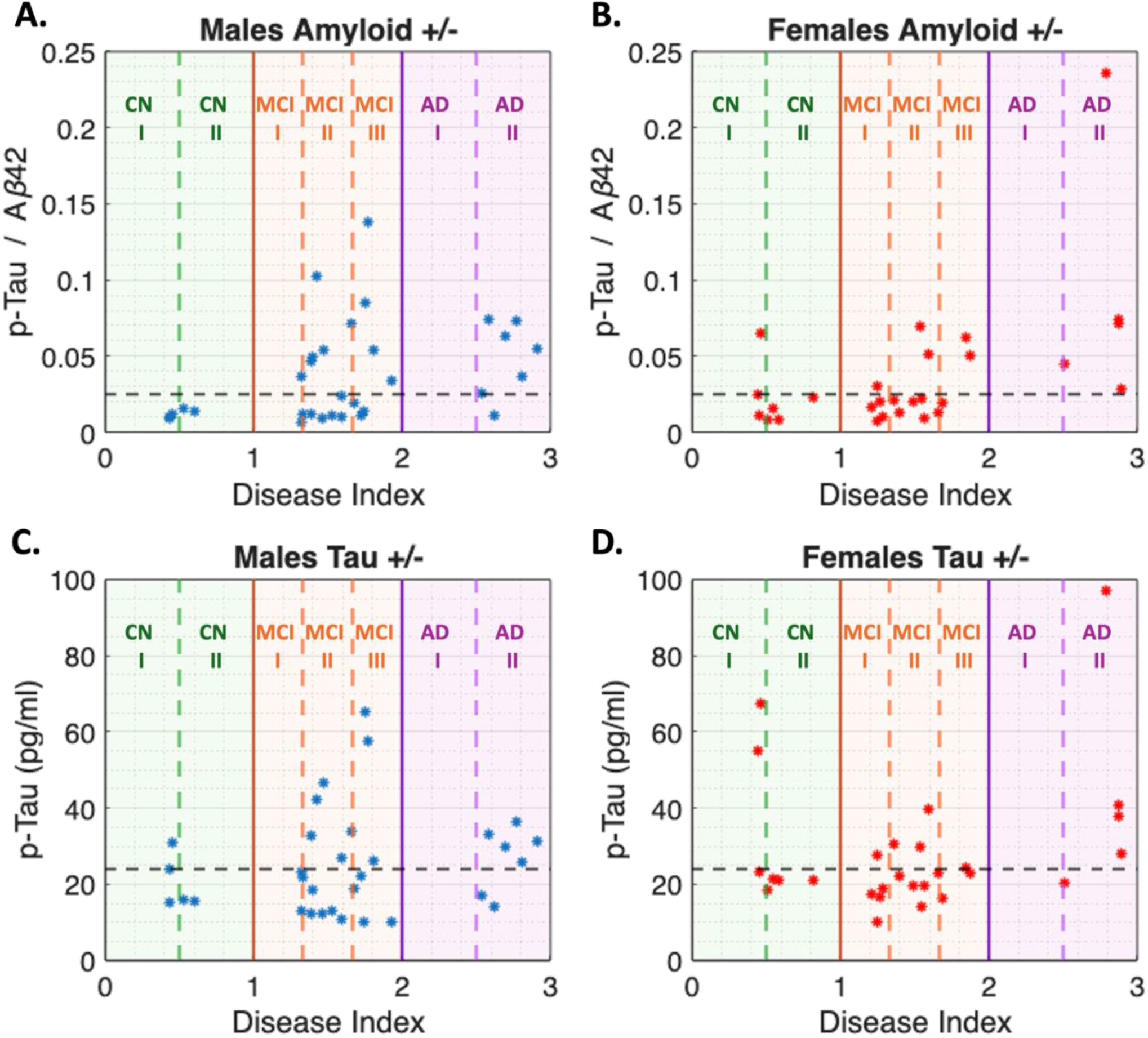
Comparison between the Disease Index and CSF Biomarkers for Amyloid-β and p-Tau. (A,B) Scatter plots display the ratio of p-Tau-181 to Aβ42 and the disease index in (A) males and (B) females. The horizontal line represents the cutoff level; values above this threshold indicate amyloid positivity. Amyloid positivity is more frequent in MCI II and higher stages, whereas subjects with MCI I or lower are predominantly amyloid-negative. (C,D) Scatter plots show CSF p-Tau-181 concentration and the disease index in (C) males and (D) females. Values above the horizontal line indicate tau-positive individuals, demonstrating that tau abnormalities also increase with higher disease index grades and emerge primarily in MCI II and subsequent stages. Both sexes exhibit similar patterns for amyloid and tau abnormalities. Importantly, the disease index was derived independently of amyloid- and tau-readouts. Shaded backgrounds denote disease stage, and dotted lines mark the boundaries between disease grades.

## 4. Discussion

This study builds upon our previous research by further elaborating on the MVD pattern^13^ (Type1 Uncoupling (T1U)⟶Type2 Uncoupling (T2U)⟶Prodromal (PC)⟶Neuro-Metabolic-Vascular Failure (NMVF)) by incorporating a third Cartesian axis derived from clinical assessments. This inclusion enables the visualization of the progression of each brain region relative to cognitive decline via a composite readout that combines multiple CCAs into a single cognitive index per brain region (Fig.1). Furthermore, these findings suggest the need for a multidimensional, multimodal approach to delineate disease etiology and improve disease staging. Our results demonstrate that although certain brain regions exhibit slower progression, the overall cluster declines across the AD spectrum. However, the extent of the spread of the group of brain regions across the cognitive degradation axis increases as the disease advances (Fig.1). For instance, in EMCI subjects, the cognitive index scores span the interval [0,1], whereas in AD subjects, this interval ranges from [4,10], with larger values indicating greater decline.

Our findings indicate that the 3D MVD pattern exhibits a spiral trajectory in key at-risk brain regions across the Cartesian space, as generated from perfusion, metabolic, and cognitive assessment readouts (Fig.3.C/Fig.S1). This observation aligns with previous preclinical and clinical studies suggesting that the brain undergoes restructuring due to bioenergetic alterations,^16-18^ synaptic pruning,^61,62^ and Excitatory/Inhibitory imbalance^63^ during AD progression. These alterations manifest clinically as memory loss, motor impairment, cognitive decline, language dysfunction, and visuospatial deficits.^16,61,63^ Consequently, these findings suggest that during phases characterized by MVD, clinical manifestations are subtle due to compensatory mechanisms in the brain aimed at preserving optimal function.^64,65^ Conversely, when the brain fails to maintain this quasi-stable bioenergetic state and transitions into prodromal and NVMF phenotypes, the clinical manifestations of AD become more evident as the pace accelerates late in disease (Fig.1.D,H), thus resulting in neurodegenerative damage caused by cerebral metabolic and/or perfusion insufficiency.^66^ These findings suggest that this approach may detect subtle changes well before amyloid or tau deposition, and may stratify subjects based on the current location of brain regions relative to the population regional trajectory.

By analyzing key brain regions related to AD, the mid-temporal gyrus exhibits a spiral curvature, with females showing a more pronounced trajectory than males (Fig.S1). This suggests that MVD alterations between sexes manifest differently, with males following the complete MVD pattern and females undergoing a faster route to more prominent clinical symptoms. This may also imply that susceptible regions between males and females may vary across key brain regions (Fig.S1), with the curvature of the trajectories suggesting differences in brain region resilience between sexes. These data are consistent with reports on the age of onset and severity of AD between the sexes, independent of *APOE* status, where women have a 56% greater likelihood of developing AD relative to men of the same age and genetic status.^67^ To validate these observations, a more detailed study would be required that tests individual trajectories, and compares these against population norms.

The observed separability of the 3D multi-domain clusters in the cross-sectional disease stage population, together with the high performance of AdaBoost as a classifier (Fig.2.A), suggests that integrating MVD signatures with a composite score derived from CCA readouts provides substantial discriminative information for data-driven staging of AD. The model’s overall AUC of 0.9479 and accuracy of 93.33%, with only 6 subjects out of 75 misclassified in the test set, indicate that these trajectories are not only descriptive but also encode robust boundaries between disease stages that can be used for individualized classification (Fig.2.B-G). Importantly, the high precision and specificity results across sexes show that the classifier is both confident and conservative, indicating a low number of false positives, which is essential for potential clinical translation. The fact that misclassifications are concentrated in the adjacent stages further suggests that the model is sensitive to biologically ambiguous, transitional phenotypes rather than making random errors, aligning with the known continuum of pathology and symptoms onset.^68^ Importantly, our machine learning approach can predict AD stage and grade in the absence of amyloid and tau with high precision and accuracy (Fig.2.F-G/Fig.5). Together, these findings position the 3D multi-domain readouts as promising candidates for data-driven model-based staging tools that may complement or refine existing biomarker frameworks for early detection and stratification in AD.

The distribution of the disease index across the AD spectrum indicates that it serves as a coherent, continuous measure of disease burden rather than a noisy aggregation of heterogeneous readouts (Fig.4.A-B). Within each clinical disease stage, the disease index approximated a Gaussian distribution, implying internally consistent grading and suggesting that individuals assigned to the same stage share a common quantitative disease profile. When the stage-specific distributions were overlaid, they showed a clear separation and an orderly shift across the disease index interval, supporting the notion that our grading framework captures a smooth transition along the AD continuum while preserving within-stage variability that may be clinically meaningful. The disease index also showed a substantially stronger correlation with disease stage than any single CCA and was more strongly associated with each CCA than the CCAs were with one another, underscoring its capacity to integrate multi-domain information into a more faithful representation of overall disease severity (Fig.4.C-D). Critically, although the index was derived independently of amyloid and tau measurements, higher grades were associated with a systematic shift from predominantly amyloid- and tau-negative profiles in early grades to near-universal amyloid and p-tau positivity in AD I and AD II, aligning the index with the biomarker-defined Alzheimer’s pathophysiologic continuum (Fig.5). This convergence between a data-driven, multimodal disease index and canonical CSF Amyloid±/Tau± profiles suggests that the proposed staging and grading framework (Fig.3.A) and scale (Fig.3.F) can bridge clinical, imaging, and molecular perspectives on AD progression, and may serve as a robust substrate for early detection, risk stratification, and therapy monitoring.

As with any research, it is important to acknowledge the limitations of this study. Firstly, the population was not stratified by age, which may be problematic, as cognitive decline is not exclusively observed in AD subjects; it may also occur in CN subjects relative to their chronological age. Secondly, the *APOE* genotype could influence the lengths of the brain region trajectories; therefore, a more comprehensive evaluation is needed to explore these factors. Third, in this study, six regions were excluded due to their relationship to the automics functional category and the lack of a surrogate measure for this; hence, a study evaluating autonomics changes using a different surrogate measure (e.g., default-mode networks) would be beneficial for including these regions. Additionally, since this study was conducted at the population level, a longitudinal study may be beneficial to elucidate and further refine the trajectories of the brain regions and their respective cubic spline fits. Fourth, evaluating the relationship between inter-stage spline lengths and regional susceptibility, and combining other CCA tests, imaging modalities, and blood/CSF biomarkers, may be beneficial for more precise AD staging. Fifth, in the grading scheme, there is an artificial truncation because the subjects’ ages in this cohort range from 55 to 95 years. It would be of interest to fit the model to younger subjects (35-55) to better approximate the distribution of CN grades. Lastly, it would be of interest to determine the minimum amount of data required by ML models to maintain a high level of accuracy, as well as to evaluate performance across alternative combinations of imaging modalities, CCA, and blood/CSF biomarkers.

In summary, our study provides a novel approach to refining disease stratification and identifying at-risk brain regions using imaging biomarkers and clinical evaluations. The integration of metabolic, perfusion, and clinical biomarkers may provide the means required for a precise medicine approach for patient monitoring, AD-stage stratification, and evaluation of therapeutic effectiveness.

## Acknowledgements

Data collection and sharing for this project was funded by the Alzheimer’s Disease Neuroimaging Initiative (ADNI) (National Institutes of Health Grant U01 AG024904) and DOD ADNI (Department of Defense award number W81XWH-12-2-0012). ADNI is funded by the National Institute on Aging, the National Institute of Biomedical Imaging and Bioengineering, and through generous contributions from the following: AbbVie, Alzheimer’s Association; Alzheimer’s Drug Discovery Foundation; Araclon Biotech; BioClinica, Inc.; Biogen; Bristol-Myers Squibb Company; CereSpir, Inc.; Cogstate; Eisai Inc.; Elan Pharmaceuticals, Inc.; Eli Lilly and Company; EuroImmun; F. Hoffmann-La Roche Ltd and its affiliated company Genentech, Inc.; Fujirebio; GE Healthcare; IXICO Ltd.; Janssen Alzheimer Immunotherapy Research & Development, LLC.; Johnson & Johnson Pharmaceutical Research & Development LLC.; Lumosity; Lundbeck; Merck & Co., Inc.; Meso Scale Diagnostics, LLC.; NeuroRx Research; Neurotrack Technologies; Novartis Pharmaceuticals Corporation; Pfizer Inc.; Piramal Imaging; Servier; Takeda Pharmaceutical Company; and Transition Therapeutics. The Canadian Institutes of Health Research is providing funds to support ADNI clinical sites in Canada. Private sector contributions are facilitated by the Foundation for the National Institutes of Health (www.fnih.org). The grantee organization is the Northern California Institute for Research and Education, and the study is coordinated by the Alzheimer’s Therapeutic Research Institute at the University of Southern California. ADNI data are disseminated by the Laboratory for Neuro Imaging at the University of Southern California. We thank Dorsey Faught from MIM Software Inc who helped with the use and development of workflows in MIM.

## Author’s Contributions

JAKCC contributed to the study design, data collection and management, data analysis and interpretation, data quality assessment, literature review, and manuscript drafting. SAC participated in study design, data collection and management, data analysis, and quality assessment. ORS did literature search for functional analysis of the brain regions. PS provided statistical and image analysis advice, participated in data interpretation, and helped write the manuscript. PRT contributed to the study design, data analysis and interpretation, and helped write the manuscript. JAKCC, PRT, and SAC had access to all data. JAKCC, SAC, PS, and PRT verified the data and results. All authors helped revise the manuscript. All authors had final responsibility for the decision to submit for publication.

## Conflict of Interest

We declare no competing interests.

## Data Sharing

The data that support the findings of this study were obtained from the Alzheimer’s Disease Neuroimaging Initiative (ADNI), which is available from the ADNI database (https://adni.loni.usc.edu) upon registration and compliance with the data use agreement.

## Ethics Committee Approval

Ethics approval was not required for this study.

## Role of Funding Source

JAKCC was supported by a T32 post-doctoral fellowship from Stark Neuroscience Research Institute (NIH grant T32AG071444). The funders had no role in study design, data collection and analysis, decision to publish, or preparation of the manuscript.

## Notes

### Competing Interest Statement

The authors have declared no competing interest.

