## Supplemental Materials for "Multidomain Analysis of Clinical Cognitive Assessments and Imaging Data in Alzheimer’s Disease Accurately Predicts Disease Stage and Grade Independent of Amyloid and Tau"

**Supplemental Material**

**Supplemental Table S1 – Single PLD ASL MRI Acquisition used in ADNI 2/3 cycles.**

| | **Siemens 2D PASL (ADNI 2/3)** | **Siemens 3D PASL (ADNI 3)** | **GE 3D pCASL (ADNI 3)** |
|---|---|---|---|
| Software Versions | 20VB17 | Prisma 20180612<br>Prisma D13<br>Prisma VE11C<br>Skyra E11<br>Skyra VE11<br>Magento Vida-XT | 25x<br>Widebore 25x |
| Repetition Time (TR) | 3400 ms | 4000 ms | 4888 ms |
| Echo Time (TE) | 12 ms | 20.26 – 21.80 ms | 10.528 |
| Field of View | 256 x 256 $mm^2$ | 240 x 240 $mm^2$ | 240 x 240 $mm^2$ |
| Acquisition Matrix | 64 x 64 | 64 x 64 | 128 x 128 |
| Reconstruction Matrix | 64 x 64 | 128 x 128 | 128 x 128 |
| Slice Thickness | 4 mm | 4.5 mm | 4 mm |
| Tag Thickness | 100 mm | N/A | N/A |
| Number of Slices | 24 | 32 | 40 |
| Bandwidth | 2368 Hz/pix | 2442 – 2604 Hz/pix | 976.6 Hz/pix |
| Background Suppression | Yes | Yes | Yes |
| M0 Available | Yes | No | Yes |
| Control-Tag Pairs | Yes, 52 | Yes, 10 | Yes, 40 |
| Mode/Readout | PICORE<br>Q2TIPS | GRASE | Stack of spirals |
| Bolus Duration | 700 ms | 800 ms | 1800 ms |
| Inversion Time / PLD | 1200 ms | 2000 ms | 2025 ms |
| Number of Repeats | 54 | 10 | 3 |
| Acquisition Time | 6:02 min | 5:24 – 8:04 min | 6 min |

### Supplemental Table S2 – FDG Acquisition and Reconstruction Protocol used in ADNI 2/3 cycles.

| Phase | Manufacturer | Model | Radiotracer (mCi) | Scan Start Time (min) | Acquisition Mode | Duration and Framing | Reconstruction Method | Method Parameters | Grid | FOV | Slice Thickness (mm) | Smoothing |
|---|---|---|---|---|---|---|---|---|---|---|---|---|
| ADNI2 | GE | Discovery STE / VCT | 4.5 – 5.5 | 30 | N/A | 30 min, six x 5min frames | 3D IR | 4 iterations; 20 subsets | 128 x 128 | 256.00 | 3.270 | None |
| | | Discovery ST | | | | | | 4 iterations; 21 subsets | | | | |
| | | Discovery RX | | | | | | | | | | |
| | | Discovery LS | | | | | 3D FORE IR | | | | 4.250 | |
| | | Advance | | | | | | | | | | |
| | Phillips | Gemini TF | | | N/A | 30 min, six x 5min frames | 3D LOR-RAMLA | N/A | 128 x 128 | 256.00 | 2.000 | Sharp |
| | | Gemini / Gemini GXL | | | | | | | | | | |
| | | Allegro | | | | | | | | | | |
| | Siemens | BioGraph mCT | | | LIST-MODE | 30 min, six x 5min frames | OSEM3D | 4 iterations; 12 subsets | 400 x 400 | 407.20 | 2.027 | None |
| | | | | | No LIST-MODE | Two 15min scans | | | | | | |
| | | BioGraph TruePoint (1093/1094) | | | LIST-MODE | 30 min, six x 5min frames | OSEM2D | 4 iterations; 14 subsets | 336 x 336 | 341.04 | 2.027 | None |
| | | | | | No LIST-MODE | Two 15min scans | | | | | | |
| | | | | | LIST-MODE | 30 min, six x 5min frames | OSEM2D | 4 iterations; 16 subsets | 336 x 336 | 341.04 | 2.027 | None |
| | | | | | No LIST-MODE | Two 15min scans | | | | | | |
| | | BioGraph HiRes (1080) | | | LIST-MODE | 30 min, six x 5min frames | OSEM2D | 4 iterations; 14 subsets | 168 x 168 | 341.21 | 2.000 | None |
| | | | | | No LIST-MODE | Two 15min scans | | | | | | |
| ADNI3 | GE | Discovery 600, 610, 690, 710 | 5 ± 10% | 30 | N/A | 30 min, six x 5min frames | 3D IR | 4 iterations; 24 subsets | 192 x 192 | 256.00 | 3.270 | None |
| | | Discovery STE | | | | | | 4 iterations; 20 subsets | 128 x 128 | | | |
| | | | | | | | | 4 iterations; 21 subsets | | | | |
| | | Discovery ST | | | | | | | | | | |
| | | Discovery LS | | | | | 3D FORE IR | | | | 4.250 | |
| | | Advance | | | | | | | | | | |
| | Phillips | Ingenuity TF | | | N/A | 30 min, six x 5min frames | BLOB-OS-TF | N/A | 128 x 128 | 256.00 | 2.000 | Sharp |
| | | Gemini TF | | | | | 3D LOR-RAMLA | | | | | |
| | | Gemini / Gemini GXL | | | | | | | | | | |
| | | Allegro | | | | | | | | | | |
| | Siemens | BioGraph mCT | | | LIST-MODE | 30 min, six x 5min frames | OSEM3D | 4 iterations; 24 subsets | 400 x 400 | 407.20 | 2.027 | None |
| | | | | | No LIST-MODE | Two 15min scans | | | | | | |
| | | BioGraph TruePoint (1093/1094) | | | LIST-MODE | 30 min, six x 5min frames | OSEM3D | 4 iterations; 21 subsets | 336 x 336 | 341.04 | 2.027 | None |
| | | | | | No LIST-MODE | Two 15min scans | | | | | | |
| | | BioGraph HiRes (1080) | | | LIST-MODE | 30 min, six x 5min frames | OSEM2D | 4 iterations; 16 subsets | 168 x 168 | 341.04 | 2.000 | None |
| | | | | | No LIST-MODE | Two 15min scans | | | | | | |

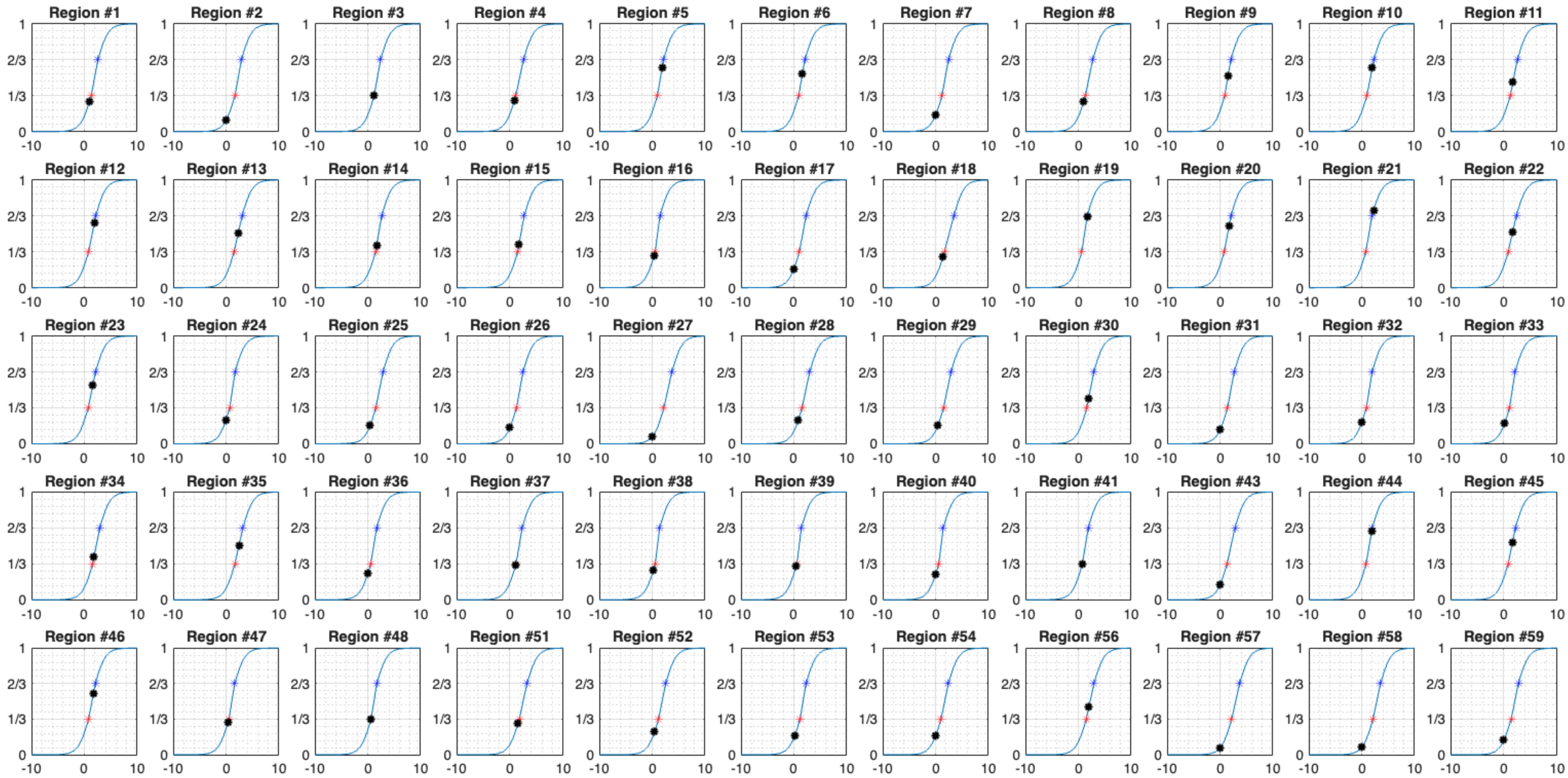


**Supplemental Figure S1 – Regional Disease Scores for a Single MCI Subject**

Regional disease scores were obtained using the disease grading scheme framework. Four regions associated with autonomic functions were omitted from this study due to the absence of suitable surrogate readouts. The figure demonstrates that each brain region progresses at distinct rates within the 3D multidomain space, as indicated by changes in regional disease scores.

### Supplemental Methods

#### A.1 Unwrapped Node Function for Optimized Length Determination

Let $C_i^3$ denote the 3D fitted cubic spline for the $i$-th brain region, constructed from population averages across five different disease stages (CN, EMCI, MCI, LMCI, AD). Each $C_i^3$ object is defined by the following parameters:

- Breaks: Knot positions where the spline changes curvature. In the UNFOLD framework, these correspond to the locations of the population averages.
- Coefficients: Polynomial coefficients for each spline segment.
- Pieces: Number of polynomials describing the spline.
- Order: Polynomial order, set to 3.
- Dimension: Data dimensionality, set to 3.

The Unwrapped Node Function for Optimized Length Determination (UNFOLD) process starts by calculating the derivative of the spline $C_i^3$ and defining the arc length function ($\mathrm{AL_i}$) as:

$$\mathrm{AL}_i(u) = \sqrt{\left(\frac{dx(u)}{du}\right)^2 + \left(\frac{dy(u)}{du}\right)^2 + \left(\frac{dz(u)}{du}\right)^2}$$

Where $x, y,$ and $z$ are the 3D coordinates of the $C_i^3$ spline. The length between two consecutive disease stages (e.g., the distance between EMCI and MCI) is then calculated by integrating $\mathrm{AL_i}$ over these stages:

$$\text{Spline Length between Stage 1 and Stage 2} = \mathrm{SL}_i(\text{Stage 1}, \text{Stage 2}) = \int_{\text{Stage 1}}^{\text{Stage 2}} \mathrm{AL_i}(u)du$$

Where $\text{Stage 1}$ and $\text{Stage 2}$ are the $C_i^3$ breaks, representing the location of the population averages. The framework then reports the total spline length, the length between consecutive stages, and the position of each population average on this linear scale.
